# Systemic application of the TRPV4 antagonist GSK2193874 induces tail vasodilation in a mouse model of thermoregulation

**DOI:** 10.1101/2021.03.12.435126

**Authors:** Fiona O’Brien, Caroline A Staunton, Richard Barrett-Jolley

## Abstract

In humans the skin is a primary thermoregulatory organ, with vasodilation leading to rapid body cooling, whereas in Rodentia the tail performs an analogous function. Many thermodetection mechanisms are likely to be involved including transient receptor potential vanilloid-type 4 (TRPV4), a widely distributed ion channel with both mechanical and thermosensitive properties. Previous studies have shown that TRPV4 can act as a vasodilator by local action in blood vessels, and in this study, we investigated whether TRPV4 activity effects *mus muscularis* tail vascular tone and thermoregulation. We measured tail blood flow by pressure plethysmography in lightly sedated *mus muscularis* (CD1 strain) at a range of ambient temperatures, with and without intraperitoneal administration of the blood brain barrier crossing TRPV4 antagonist GSK2193874. We also measured heart rate and blood pressure with and without GSK2193874. As expected for a thermoregulatory organ, we found that tail blood flow increased with temperature. However, unexpectedly we found that the TRPV4 antagonist GSK2193874 increased tail blood flow at all temperatures, and we observed changes in heart rate variability. Since TRPV4 activation stimulates the relaxation of peripheral resistance arteries (vasodilation) that would increase tail blood flow, these data suggest that increases in tail blood flow resulting from the TRPV4 antagonist may arise from a site other than the blood vessels themselves, perhaps in central cardiovascular control centres such as the hypothalamus.

## Introduction

Thermoregulation is one of the defining homeostatic processes common to mammals; core body and brain temperatures are well maintained despite challenges such as changing ambient temperature and exercise to the degree that brain temperature rarely changes outside of a 3 °C range [1-3]. Mammals detect temperatures at both central and peripheral sites and responses to changing temperatures can result both from local responses and central, hypothalamus-co-ordinated autonomic responses [4-6]. Typical thermogenic effector mechanisms include liver thermogenesis and skeletal muscle shivering whereas cooling mechanisms including behavioural changes and redistribution of blood from core to peripheral vessels [4, 5, 7]. Rodents use basal metabolic rate and non-shivering thermogenesis as their principle mechanisms for heat production, mainly because of their small size [8]. In terms of heat loss, transfer of excess heat to the environment is facilitated by so-called heat transfer zones, which are usually found at the body extremities, for example, in humans, typically, acute heat loss is mediated by redistributing blood to cutaneous vascular beds [5]. The location of critical heat transfer zones are somewhat species specific, so for example, the ear for elephants and rabbits [9, 10], head vasculature in large dinosaurs [11, 12] and the feet [13] and tail for rodents [14-16]. The tail of rodents is ideal as a heat transfer zone due to its glabrous nature [16]. It thought that vasoconstriction rather than counter-current heat exchange provides the major barrier to core-to-tail heat flow [17].

In this work, we have investigated the role of *mus muscularis* TRPV4 in this homeostatic system using a potent and selective TRPV4 inhibitor, GSK2193874. TRPV4 is one of several temperature sensitive ion channels and expressed in both the hypothalamus and the vasculature, in both smooth muscle and endothelial cells. Recently, there has been considerable interest in the immune, neuromodulatory, cardiovascular and thermoregulatory potential of small molecule TRPV4 modulatory drugs, such as GSK2193874 and HC-067047 [18-25].

TRPV4 is a relatively non-selective Ca^2+^ channel (PCa/PNa 6-10) that was first characterised as mechanosensory [26, 27], however, it is also activated by temperatures >30 °C, and so, at physiological temperatures, it would be expected to be constitutively active under basal conditions [28-30]. Activation of TRPV4 leads to vasodilation [31-34] and logically, therefore, transgenic elimination of TRPV4 (TRPV4-/-knock out) would be expected to increase blood pressure, but it does not [18, 31].

The precise contribution of TRPV4 to thermosensing and thermoregulation *in vivo* remain unclear. No changes in escape latency from heat stimuli were observed in the hotplate challenge [35, 36]. However, post subcutaneous injection of capsaicin or carrageenan, TRPV4-/-mice showed longer escape latencies from the hot surface compared to wild-type [36]. In another study, it was shown that TRPV4 is required for normal thermal responsiveness *in vivo*; on a thermal gradient, TRPV4-/-mice selected warmer floor temperatures. In addition, TRPV4-/-mice also exhibited prolonged withdrawal latencies during acute tail heating [37].

In terms of pharmacological manipulations, activation of TRPV4 with topological RN1747 decreased core temperature of *rattus norvegicus* and increased tail vasodilatation [38]. The effects of a TRPV4 inhibitor (HC067047), in the same study were mixed with increases of core body temperature with ambients of 26 and 30 C, but not 22 and 32 C.

In this study we had aimed to investigate whether the small molecule TRPV4 inhibitor, GSK2193874, would decrease tail vasodilation response to elevated ambient temperatures. As a surrogate for tail vasodilation we used tail blood flow measured by volume plethysmography [39]. We also investigated frequency domain heart rate variability (HRV). HRV is a sensitive tool that assesses the time difference between consecutive heart beats to evaluate autonomic nervous system modulation [40, 41]. Accumulating data suggests that ultra-short-range HRV can be successfully derived from as low as 30s of human ECG [42, 43] and pulse rate variability (estimation of variation in heart rate from *photo*plethysmography) has recently been successfully measured from the rat tail [44]. Potentially, measurement of HRV from tail-cuffs would be a useful 3Rs advancement, since surgery is not required. Therefore, we sought to, for the first time, (a) establish, empirically, the length of heart rate (HR) record necessary for HRV in mice and (b) perform HRV from mouse tail volume plethysmography using the CODA apparatus. HRV reflects homeostasis in thermoregulation and blood pressure control and has been shown to be modulated by thermal stimuli in humans [45].

Surprisingly, we found the TRPV4 inhibitor increased tail blood flow when measured above mouse thermoneutrality, and we saw temperature dependent changes in ultra-short-range HRV raising the possibility that TRPV4 ion channels expressed outside of the vasculature, for example in the central nervous system, may also be involved with rodent thermoregulation.

## Methods

### Animals

Fourteen female adult CD1-mice (Charles River, UK) were used. All experimental procedures were ethically approved by the University’s Animal Welfare Committee and performed under a UK Home Office Scientific Procedures licence (70/8746).

### Volume pressure plethysmography (VPR) recording

We used the CODA tail volume plethysmography (VPR) system (Kent Scientific, Torrington, CT, USA) on control CD1-mice and mice that had received the selective TRPV4 antagonist GSK2193874. Full details of warming methodology and VPR methods are included in the supplementary materials. Note, all temperatures reported are ambient temperatures read from the thermocouple.

### Statistical Analyses

Blood pressure (MAP), heart rate (HR) and blood flow statistical comparisons were made with the nlme package in R, which incorporates a repeated measures design. For HRV statistical comparisons, we used MANOVA in Minitab (PA, USA). p≤0.05 was taken as significant.

### Drugs

Midazolam was supplied by our animal service unit, but GSK2193874 (300 µg/kg, *i*.*p*.) and DMSO were obtained from Sigma-Aldrich. GSK2193874 was dissolved in DMSO at 20 mg/ml stock then diluted 1:100 before i.p. injection (0.2 mg/ml), following [23, 34]. “Control” includes 1% DMSO and volume of injection was dependent upon animal weight.

## Results

We measured MAP, HR and blood flow (Flow) in 14 animals with and without GSK2193874 over the ambient temperature range of 31ºC to 36ºC. These are plotted in two-factor (treatment, temperature) format and analysed with a repeated-measures, mixed effects design. There was a statistically significant effect of temperature on all parameters measured, MAP (**Figure 1A:** Temperature F=5.34, p≤0.05; Drug F=0.38 p>0.05, Drug x Temperature F=0.17, p>0.05), HR (**Figure 1B:** Temperature F=7.37, p≤0.05; drug F=0.68 p>0.05, Drug x Temperature F=0.23, p>0.05) and tail blood flow (**Figure 1C:** Temperature F=13.21, p≤0.005; drug F=5.57, p≤0.05, Drug x Temperature F=14.00, p≤0.0005). In the cases of HR and MAP there was no significant effect of treatment with the TRPV4 antagonist (GSK2193874). However, with tail blood flow there was both a significant increase with GSK2193874 treatment and a very highly significant interaction between temperature and GSK2193874 treatment.

**Figure 1.**
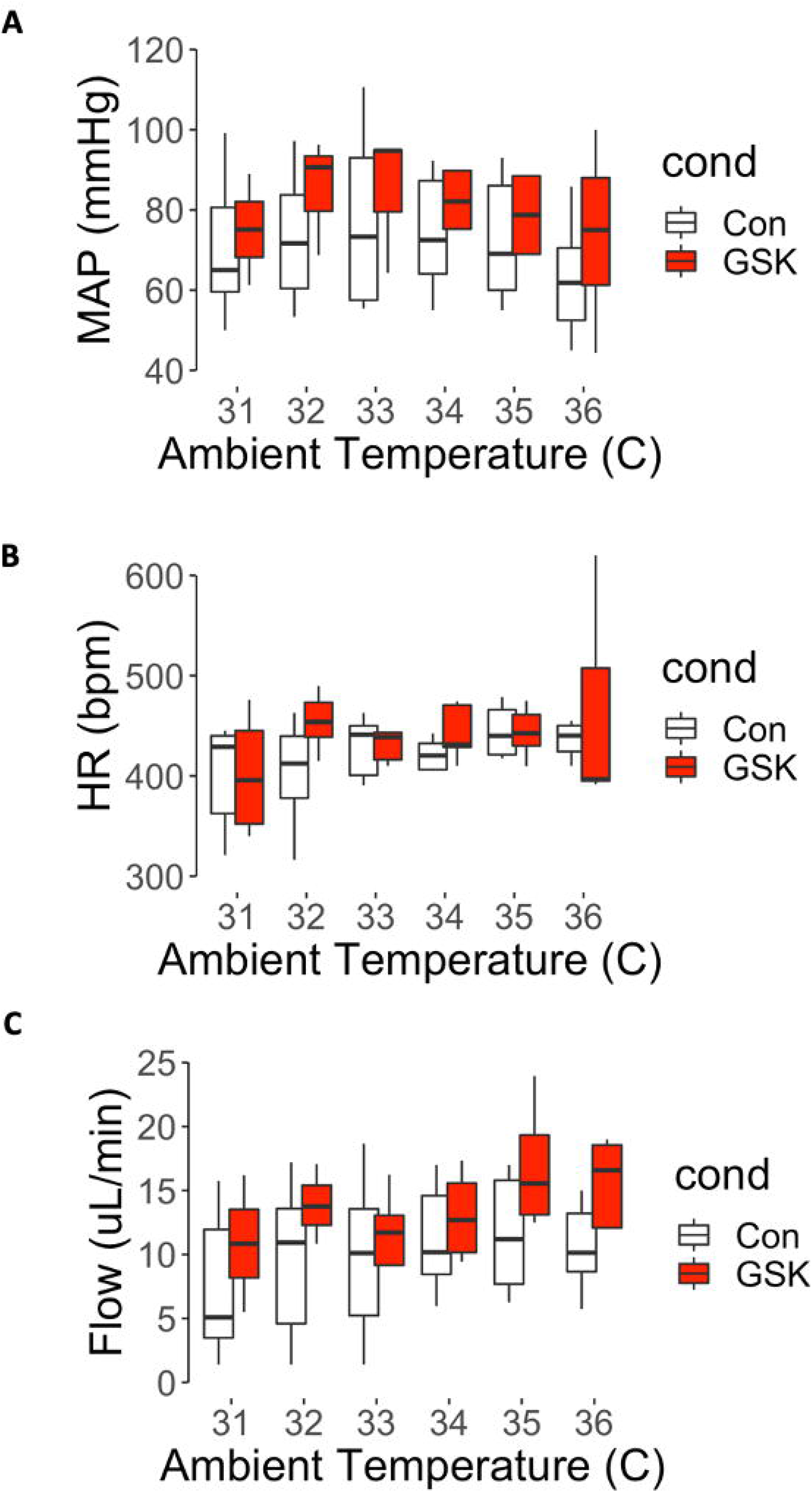
Effects of Pharmacological inhibition of TRPV4 on blood pressure, heart rate and blood flow. **(A)** Mean arterial pressure at a range of ambient temperatures in control and GSK2193874, there was a significant association with temperature, but not drug. There was also no significant association between temperature and drug (Mixed effects model: See main text for details). **(B)** Mean heart rate across a range of ambient temperature in control and GSK2193874, over-all there was a significant change of heart rate with temperature, but no significant difference with drug and no significant association between drug and temperature (Mixed effects model: See main text for details). **(C)** Tail blood flow across a range of ambient temperature in control and GSK2193874. Overall, there was a statistically significant increase of blood flow with temperature and with GSK2193874 and a significant interaction (Mixed effects model: See main text for details). Overall, *n* = 14 animals or for each temperature; 31ºC *n*= (9,4) 32ºC *n*= (10,4), 33ºC *n*= (11,6), 34ºC *n*= (10,7), 35ºC *n=* (11,7) and 35ºC *n*= (12,5). To investigate specific temperature points that were different to the 31ºC value, we treated temperature as a factor and ran the estimated-marginal means method with the R-package Emmeans. This consists of 66 pairwise comparisons and we used Benjamini-Hochberg multiple comparison correction. In control, no individual flow is significantly greater than that at 31ºC however, in GSK2193874, blood flow at both 35ºC and 36ºC was significantly greater than at 31ºC (p<0.05).

Since we were able to derive beat-by-beat heart rate records for several seconds (for example Supplementary Figure 1), we investigated whether HRV could be captured over such short periods. To test whether this was feasible, we simulated mouse heart rate interval records of decreasing length using a modified version of McSharry *et al*., 2003 [46] and then measured HRV spectral powers by the Lomb-Scargle method [47, 48] over 3000 simulations. **Figure 2A** shows that just a few seconds of ECG are sufficient to obtain a picture of the heart rate variability in a mouse, in so far as, increasing the simulation duration beyond this does not greatly affect the HRV spectra. We therefore measured HRV power in the 0.1 to 1.9Hz bands in our samples of control and GSK2193874 records (**Figure 2B, C**) and compared these statistically with a MANOVA model, over a range of temperatures. There was no over-all statistical difference with temperature, however there was a statistically different set of spectra between control and GSK2193874 treated spectra. Furthermore, with univariate analyses, there was a significant difference between treatment and control at each individual frequency except the 0.5Hz banding.

**Figure 2.**
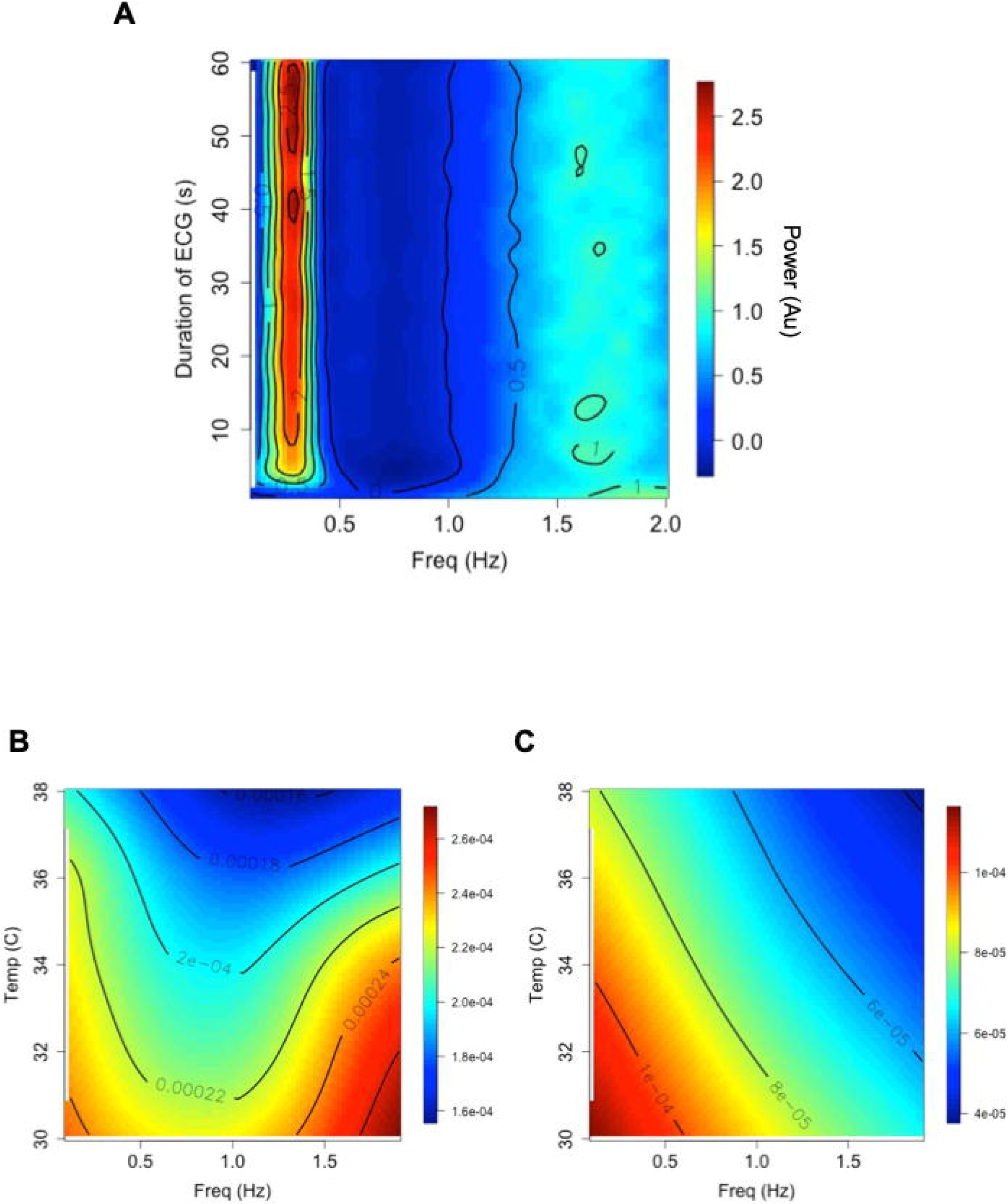
Short range heart rate variability analysis. **(A)** Stacked Lomb periodograms for 3000 simulated ECG inter-beat interval datasets. Y-axis is the duration of the simulated ECG record, X-axis is the frequency component of each Lomb-Scargle. The periodograms have been normalized and scaled therefore the power (colour bar on the right) is in arbitrary units. Below are periodogram surfaces recorded at different temperatures under control **(B)**, or after injection GSK2193874 **(C)**. Power given in the scale-bars. MANOVA (Minitab) analyses show these two distributions are significantly different, Wilks lambda p<0.05. Overall, *n* = 14 animals.

## Discussion

In this work we investigate the role of TRPV4 in rat tail blood flow with a systemic inhibitor of TRPV4, GSK2193874. Surprisingly, we find that tail blood flow is increased by GSK2193874. Inspection of Figure 1C., suggests that there is little effect of temperature, in control conditions, above (≥) 32ºC, however the largest numerical increase in flow occurred above 35ºC ambient. Possibly suggesting that this quite high ambient temperature was necessary to see the changes in TRPV4 activity. There was a detectable effect also on HRV, but no significant change in blood pressure or HR themselves.

### Blood flow, HR and MAP effects

GSK2193874 is a small lipid soluble inhibitor of TRPV4 [19] that crosses the blood-brain barrier well (brain:plasma ratio = 0.6, personal communication with Dr David Behm of GSK) and so there are several locations at which TRPV4 could potentially influence the control of blood flow in response to elevated temperatures. A non-exhaustive list of possible sites of action could include the vasculature or cardiovascular control neurones.

TRPV4 is expressed in both vascular smooth muscle and the endothelial cell lining [49]. Activation of these channels leads to clear vasodilatation. Whilst the mechanism of this vasodilatation is complex, involving both endothelial and smooth muscle cells, potential release of EDRF/EDHF and, ultimately, small local increases of Ca activate potassium channels which hyperpolarize the muscle cells and allow relaxation/vasodilatation [31-33]. A TRPV4 inhibitor would therefore be expected to cause vasodilation (or have no effect if there was no constitutive TRPV4 activity) and so it seems unlikely the increase in tail blood flow we report in this study results from direct action on the vasculature. Furthermore, if the effect of GSK2193874 were primarily on blood vessels to cause dilation, we would have expected to see an over-all drop in MAP, and possibly then a reflex increase in HR since the baroreceptor loop features in established mechanisms of cardiovascular control as well as, specifically, thermoregulation [50, 51]. We saw no change in blood pressure or heart rate, although multivariate analysis detected a small change in short-range HRV analysis. The potential for us to have missed such a baroreceptor mediated effect due Type II errors is discussed in the *limitations* section below.

A second location of TRPV4 channels that may be of relevance is the central nervous system, for example the hypothalamus [52]. It is known that other transient receptor potential channels influence the cardiovascular system via changes in sympathetic activity [53, 54]. Our own work shows that TRPV4 channels are located on pre-autonomic neurones of the hypothalamic paraventricular nucleus (PVN) and can influence cardiovascular control in response to osmotic challenge [55, 56] and this effect was abolished with a TRPV4 inhibitor [55]. At the neuronal level, we have shown that the action current frequency of parvocellular PVN neurones is dramatically reduced when TRPV4 channels are inhibited [56]. To date, there have been no studies that have explored thermoregulatory roles for TRPV4 in central cardiovascular control neurones.

### HRV effects

HRV analysis is an increasingly common method for cardiovascular assessment. In humans, for example, decreased HRV (i.e., a very steady pulse) is an independent predictor of cardiac mortality [57]. In animals too it is proving increasingly useful in a range of contexts including phenotyping transgenic animals [58], investigating cardiovascular effects of drugs [59] and predicting arrythmias [60]. Whilst there are many papers analysing HRV in mice using radiotelemetry [61] we investigated here whether it was possible do this with VPR. It has previously been shown that relatively long *photo*plethysmography recordings could be used for HRV, with high accuracy, but the present study is the first to systematically analyse how long a recording needs to be. The derivation of this short-range HRV from non-invasive apparatus may prove a useful advance in 3Rs. Since the average mouse heart rate is some 8 times that of a human, an 8s segment would be equivalent to the standard 1 minute of recording necessary to detect higher frequency components of human ECG [41]. Here, simulation shows that periodograms from very short segments of ECG are similar to that of conventional 1-minute records (Figure 2A), and these data themselves and this approach may be of field interest. In terms of the response to temperature, we did not see an overall effect on HRV, probably because temperature typically effects the low frequency powers, beyond the scope of ultra-short-range recording [45], however, GSK2193874 did significantly alter overall frequency power.

### Limitations

We measured only ambient temperature and not core temperature. We felt that the loss of this important information was necessary to avoid the disturbance of rectal thermocouple of mice in the non-invasive recording equipment. Also, we used sedation that could influence the whole animal responses and only *female mice*, unlike many males only studies [23, 34]. Furthermore, to keep the study manageable, we opted for a one antagonist dose study rather than a full in vivo dose-response curve, which would have been useful. It is difficult to predict accurately the local concentration that an ion channel will “see” when a drug given systemically will reach, but if we assume that GSK2193874 has a typical volume of distribution of between 1 and 10L/kg, our 300µg/kg dose would translate to approximately 40 to 400nM, in the order of the maximal dose for GSK2193874 on TRPV4 channels [23]. Although GSK2193874 is highly selective for TRPV4 compared to the other 200+ proteins it has been assayed against [23], repeating our studies with TRPV4-/-lines [31] would be the only way to confirm with certainty that the true target was indeed TRPV4. We encountered technical challenges too, e.g., recording VPR data below 30ºC (ambient) was unreliable, so we report a relatively limited temperature range rather than strictly hot vs cold. These limitations could be addressed by a telemetric study, but large motivation for our current approach was to utilise a non-invasive blood pressure design, for 3Rs ethical reasons. Furthermore, as in many physiological studies, statistical power was an issue. Our initial design (see supplementary information) included a power analysis for heart rate and blood pressure; which made a number of assumptions but passed 80% power with around 8 or 9 animals. We then used 14, however, we were not able to get all conditions for all animals and so the final statistical power could be below 80%. We have hypothesised that an increase of blood flow, by TRPV4 antagonist in the absence of significant changes in MAP/HR would be compatible with a central mechanism of vasodilation. However, if we simply missed changes due to a type II error, the vasodilation could result from baroreflex-mediated mechanisms. This could be addressed by either increasing animal numbers or by repeating similar experiments with surgical or pharmacological block of the baroreceptor reflex [62].

In conclusion, this whole animal study shows that a TRPV4 antagonist has a significant effect on tail blood flow, in the context of thermoregulation, but its site of action, and the mechanism of such modulation remain to be determined. We also demonstrate non-invasive measurement of frequency domain HRV analysis from very short-range data that may prove useful in future 3Rs friendly research.

## Supporting information

power analysis HR

power analysis BP

## Acknowledgements

This project received funding from BBSRC: BB/T002115/1, BB/T002115/1

BB/S008136/1 AND BB/N003020/1 awarded to RBJ.

