## Supplementary material for "Systemic application of the TRPV4 antagonist GSK2193874 induces tail vasodilation in a mouse model of thermoregulation": supp info 1

**Supplementary Materials**

**Extended Methods**

*Volume pressure recording (VPR) plethysmography*

A representative example of the method (Supplementary Figure 1A), which is sophisticated plethysmography, using microsensors. An occlusion cuff is inflated to 250 mmHg. This halts blood flow to the tail, and the tail volume reaches a minimum the tail volume is detected/estimated by the “VPR pressure sensor”. Then the tail-cuff is deflated over 15 seconds. As pressure in this tail-cuff pressure decreases over this time there comes a point at which blood pressure exceeds the tail-cuff pressure and blood can begin to flow into the tail. This is detected by the VPR pressure sensor. From such data, several features can be extracted by the CODA software such as blood pressure, tail blood flow, tail volume and heart rate. Supplementary Figure 1B shows an expanded view of the data in Supplementary Figure 1A, and Supplementary Figure 1C shows how with sufficient amplification and filtering individual pulse can be extracted, and peaks detected to allow heart rate measurement, independently of the CODA software.

To measure heart rate as accurately as possible, raw cuff pressure traces were exported as csv files and analysed in MATLAB (See Figure 1C). Here, they were digitally filtered and peaks detected with MATLAB’s peak detector function.

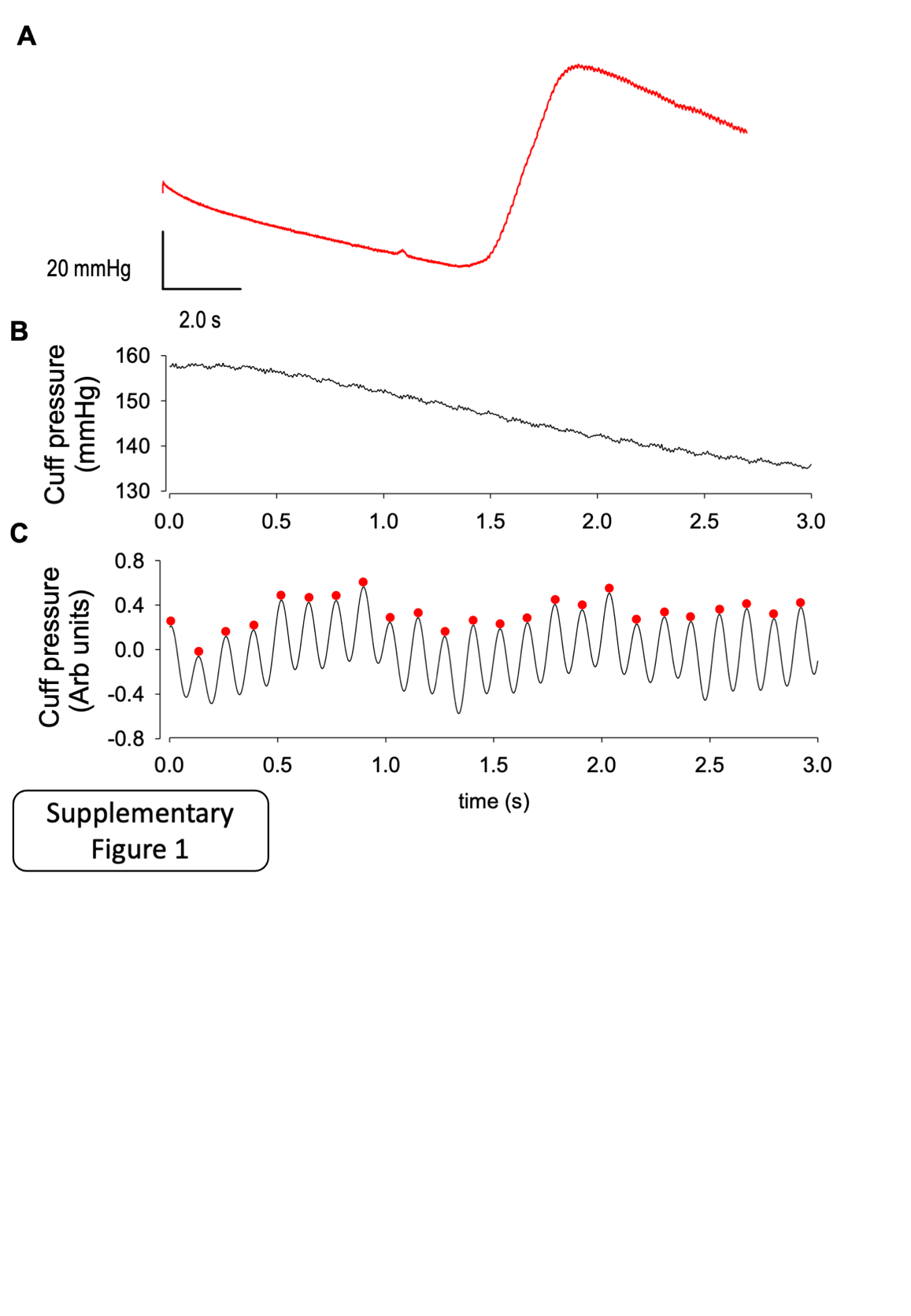

**Supplementary Figure 1. Pharmacological inhibition of TRPV4 increases tail blood flow in CD1 mice**

(A) Representative examples of complete (single) VPR. The system comprises two tail cuffs, an occlusion cuff and a VPR-sensor cuff. The occlusion cuff is placed at the base of the tail and the VPR sensor cuff over the middle of the tail. Initially the occasion cuff is inflated to approximately 250mmHg. This prevents blood entering the tail and the pressure in the VPR sensor steadily decreases. Once the occlusion cuff drops sufficiently blood returns to the tail and causes activation of the VPR sensor. This is factory calibrated to allow calculation of blood flow/volume. The wave/peak on the right-hand side is where the tail volume increased. Following the VPR pressure-peak, the pulse pressure can be seen in the VPR-pressure traces. An enlarged view is shown in (B). (C) This section of recording can then be filtered and amplified to allow each pulse pressure peak to be identified (filled red circles) and further analyzed. This can be automatically by the CODA software, but this figure was created via a MATLAB script that allows more detailed heart rate analysis.

*Temperature Challenges*

Ambient temperature was increased slowly using heating pads under their housing and insulating the cage lids so the entire animal is exposed to the new temperature. Ambient temperature changed at a rate of approximately 0.25°C per minute. Temperature was monitored by thermocouple place alongside the animal in the enclosure during the experiment whilst the CODA device was continuously recording. All temperatures reported are ambient temperatures read from the thermocouple. Core temperature was not recorded since rectal probe insertion disturbed the mouse subjects. Each animal was exposed to as many different temperatures as possible to add within-subject statistical power. Some animals received control followed by GSK2193874, but some animals were given just GSK2193874 because we could not be sure when it would have left the animal system and follow test with control. This design is all encoded in the statistical analyses described below.
