## Supplementary material for "Systemic application of the TRPV4 antagonist GSK2193874 induces tail vasodilation in a mouse model of thermoregulation": power analysis HR

### R Notebook

powerSim: estimates power by simulation Performs a power analysis for a mixed model powerSim(fit, test = fixed(getDefaultXname(fit)), sim = fit, fitOpts = list(), testOpts = list(), simOpts = list(), seed, …) fit=fitted model object test=specifies test to perform sim=object to simulate from Tutorial available here: https://humburg.github.io/Power-Analysis/simr_power_analysis.html

```
## Loading required package: lme4
```

```
## Loading required package: Matrix
```

```
##
## Attaching package: 'simr'
```

```
## The following object is masked from 'package:lme4':
##
##     getData
```

#Create covariates Start by setting up a data frame with 12 subjects, 2 drugs classes and 10 temperatures, with and without vehicle

```
## [1] 528
```

```
## [1] 528
```

```
## [1] 528
```

```
## [1] 528
```

```
## [1] 528
```

##Specify the parameters for the model. We'll use the model Structuring witn all the possible interactions would be inpractical, so keep simple. HR~temp+drug+veh+(1|subject)+error Parameters should come from literature

```
## [1] 13
```

##Create model The makeLmer function from the simr package allows us to combine all this information to create a fitted lmer model from scratch.

```
## Linear mixed model fit by REML ['lmerMod']
## Formula: y ~ Drug + Temp + Veh + (1 | ID)
##    Data: covars
## REML criterion at convergence: -55.5127
## Random effects:
##  Groups   Name        Std.Dev.
##  ID       (Intercept) 0.4472
##  Residual             0.2000
## Number of obs: 528, groups:  ID, 12
## Fixed Effects:
## (Intercept)     DrugNone       Temp23       Temp24       Temp25       Temp26
##      600.00         0.15         0.15         0.15         0.15         0.15
##      Temp27       Temp28       Temp29       Temp30       Temp31       Temp32
##        0.15         0.15         0.15         0.15         0.15         0.15
##      VehH20
##        0.01
```

##Power analysis Once you have a fitted lmer model, whether it was fitted to real data or created from scratch, you can use that to simulate new data and assess the required sample size.

```
## Power for model comparison, (95% confidence interval):
##       100.0% (69.15, 100.0)
##
## Test: Likelihood ratio
##       Comparison to y ~ Temp + [re]
##
## Based on 10 simulations, (0 warnings, 0 errors)
## alpha = 0.05, nrow = 528
##
## Time elapsed: 0 h 0 m 0 s
```

We can test for the effect of time in the same way.

```
## Power for model comparison, (95% confidence interval):
##       100.0% (69.15, 100.0)
##
## Test: Likelihood ratio
##       Comparison to y ~ Drug + [re]
##
## Based on 10 simulations, (0 warnings, 0 errors)
## alpha = 0.05, nrow = 528
##
## Time elapsed: 0 h 0 m 0 s
```

```
## boundary (singular) fit: see ?isSingular
## boundary (singular) fit: see ?isSingular
## boundary (singular) fit: see ?isSingular
## boundary (singular) fit: see ?isSingular
```

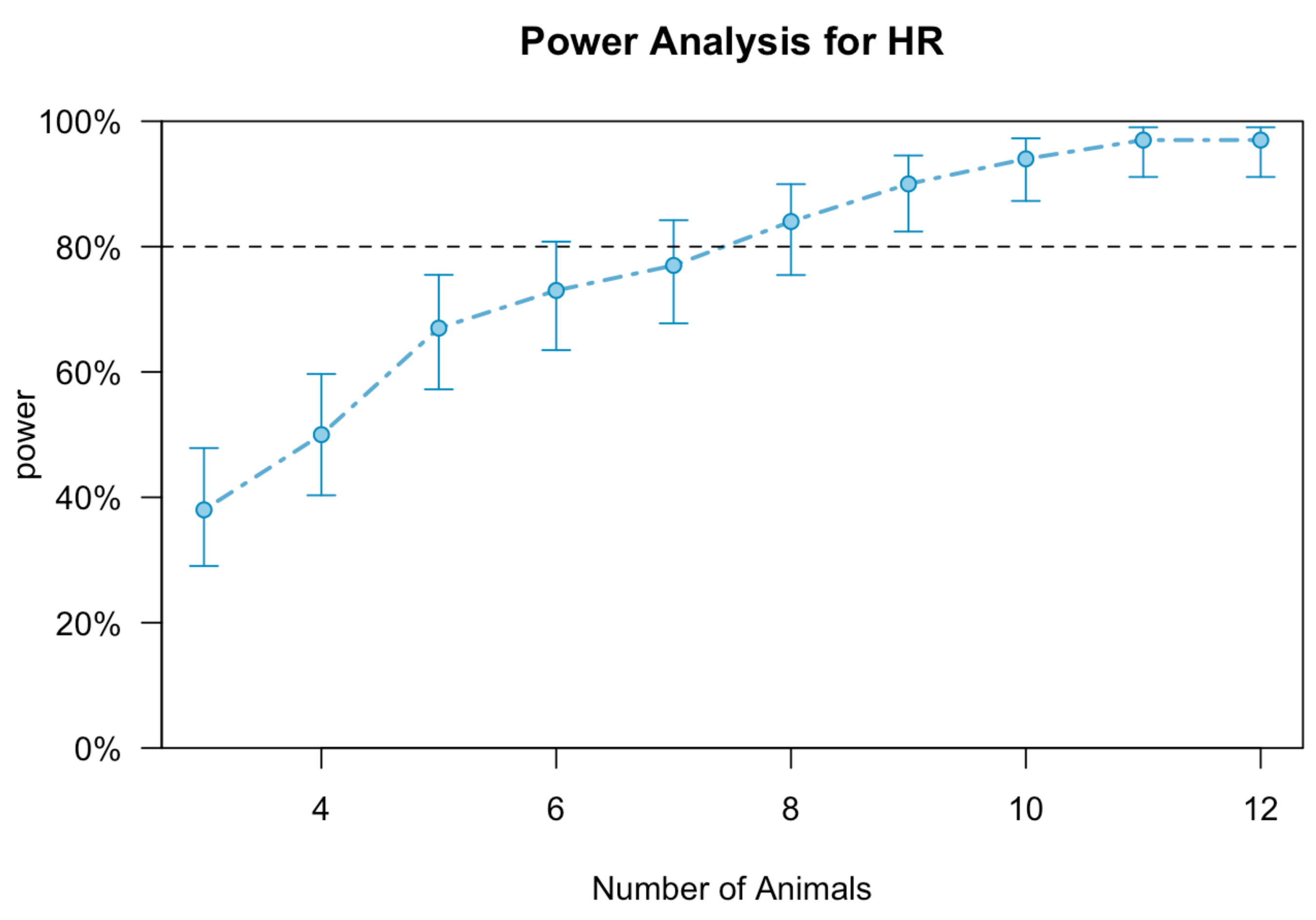


```
## integer(0)
```
